# Genotypic Responses of Rice to Ammonium and Nitrate under Contrasting Water Regimes: Agronomic, Biochemical, and Gene-Expression Analyses

**DOI:** 10.64898/2026.10.04.756527

**Authors:** Helan Baby Thomas, Toshy Agarwal, Satish Verulkar

## Abstract

Considering the increasing water and labor constraints in rice production, direct-seeded rice is becoming increasingly important. However, direct-seeded rice is vulnerable to water stress and inefficient nutrient use. Nitrogen use efficiency in rice is generally low, and improving nitrogen acquisition and assimilation is important for sustainable productivity. In this study, 48 recombinant inbred lines were evaluated under ammonium and nitrate nutrition in irrigated and rainfed environments across two cropping seasons. Agronomic, physiological, biochemical and molecular responses were assessed. Significant effects of genotype, nitrogen form, environments and their interactions were observed for several traits. Rainfed conditions reduced grain and biological yield compared with irrigated conditions. Ammonium produced a higher mean grain yield than nitrate by approximately 26.2 g m^-2^ under irrigated conditions and 154.2 g m^-2^ under rainfed conditions, although the responses varied among genotypes. Biological yield and SPAD values were generally higher with ammonium, whereas total chlorophyll showed season and environment-dependent responses. Glutamine synthetase activity was highest under ammonium treatment, while glutamate synthase activity was higher under nitrate treatment. Grain yield showed strong positive correlations with biological yield, but correlations between grain yield and GS or GOGAT activity were not significant. Expression of OsGln1.2, OsGln2 and OsGlt1 varied according to genotype, tissue and nitrogen treatment. Overall, the results demonstrate that responses to nitrogen forms are genotype- and environment-dependent, and that ammonium may provide a yield advantage under rainfed conditions. Integrating field performance with biochemical and molecular traits may help identify rice genotypes suited to water-limited systems.

## 1. Introduction

Rice has sustained over than half of the world’s population for ages, with global consumption reaching about 530.239 million metric tonnes in the 2024-25 crop year (STATISTA 2025). Rice is the third most cultivated cereal globally, with a production of 522.62 million metric tonnes (USDA 2024). It is predominantly cultivated in Asia, where India has the highest acreage, producing 137.83 million metric tonnes (USDA 2024). The two major subspecies, indica and japonica, account for more than 80% of the world’s rice cultivation (Cai, Liu et al. 2007). Among the rice-growing states in India, Chhattisgarh, a central eastern state also known as the ’Rice Bowl of India,’ produced 9.7 million metric tonnes in 2024 (Outlook 2024). In the current scenario, it is imperative to enhance global rice production to meet the growing food demands driven by population growth. Increasing arable land might not be a feasible solution, but identifying nutrient-efficient genotypes can increase productivity and annual yield (Vinod and Heuer 2012). Rice production requires a large amount of water, with 75% of global paddy cultivation relying on irrigation systems. Rice production is severely affected by various biotic and abiotic stresses, with drought being a major constraint in regions where irrigation sources are limited and dependent on rainfall (Fahad, Adnan et al. 2019, Sarma, Kashtoh et al. 2023). Drought interferes with nutrient acquisition by the roots and their mobilization into sink tissues, reducing nutrient concentration in plant tissues. Water resources are dwindling daily due to climate change, groundwater depletion, pollution, and other environmental changes (Mishra 2023, Davamani, John et al. 2024). Consequently, the area under direct-seeded rice is gradually increasing, as it requires less water than lowland rice and may eventually cover the majority of paddy-cultivated areas (Ishfaq, Akbar et al. 2020). Under direct seeding, the anaerobic conditions are changed into aerobic, which would further lead to the change in the absorption of different forms of nutrients.

Nitrogen is one of the most limiting nutrients in agricultural systems as substantial losses can occur through leaching, volatilization and denitrification. (Hirel, Le Gouis et al. 2007) reported that the two-fold increase in agricultural food production in the world was associated with a seven-fold increase in the use of nitrogenous fertilizers. Nitrogen is critical for achieving optimum yield in lowland rice (Singh 2006). Consequently, the application of nitrogen (N) fertilizer has become a decisive factor in attaining high yields in rice. However, the nitrogen use efficiency (NUE) of rice is only 33%, the lowest among cereals. The unutilized nitrogen leaches into water bodies, causing pollution, affecting health, and damaging biodiversity. Nitrogen availability is reported to be impacted by soil water scarcity (Gonzalez-Dugo, Durand et al. 2010, Flynn, Comas et al. 2023). (Smika, Haas et al. 1965) observed that nitrogenous fertilizers are not utilized efficiently in the absence of sufficient water. Therefore, the amount of nitrogen available to plants can be improved by managing the interaction between fertilizer, soil, and water and by harnessing the innate efficiency of genotypes to utilize the available nitrogen and achieve high productivity.

Rice has an inherent ability to utilize ammonium and nitrate forms, although ammonium is generally preferred over nitrate as most of the rice around the world is grown under irrigated conditions (Chen, Li et al. 2024, Guo, Zhang et al. 2025). The rice roots and the whole metabolic system of the plant have evolved and adapted for the efficient utilization of ammoniacal form of nitrogen, while the nitrate form is preferred during seedling and early vegetative phases. Nitrogen assimilation in the plant is controlled by many gene families which are involved in NH_4_^+^ and NO_3_^-^ uptake, translocation, utilization, etc. Several membrane transporters have been identified as key mediators in the uptake of both inorganic nitrogen forms (NH_4_^+^ and NO_3_^-^) and organic nitrogen compounds from the soil into plant roots (Shinozaki, Yamaguchi-Shinozaki et al. 2003, Nazish, Arshad et al. 2022, Guo, Zhang et al. 2025). Following uptake, nitrate (NO_3_^-^) is reduced to ammonium (NH_4_^+^) via the coordinated action of the nitrate transporter system and the enzymes nitrate reductase (NR) and nitrite reductase (NiR). The resulting ammonium, derived either from nitrate reduction or direct absorption, is subsequently assimilated into amino acids within plastids through the glutamine synthetase/glutamate synthase (GS/GOGAT) cycle (Campbell 2002, Lea and Miflin 2010).

Glutamine synthetase (GS) plays a central role in nitrogen metabolism, particularly in the assimilation of ammonium and mitigation of its potential toxicity (Fortunato, Nigro et al. 2023). Its significance in enhancing nitrogen use efficiency (NUE) has been demonstrated in rice (Wu, Zhang et al. 2021). Overexpression of cytosolic GS1 (OsGS1.1) and chloroplastic GS2 (OsGS1) in transgenic rice has been shown to improve tolerance to osmotic salinity stress during the seedling stage (Cai, Zhou et al. 2009, James, Borphukan et al. 2018). However, comparatively few studies have evaluated genetically diverse rice lines under ammonium and nitrate nutrition across both irrigated and rainfed environments while integrating agronomic, physiological, biochemical and molecular traits.

Therefore, the present study aimed to characterize 48 rice recombinant inbred lines under ammonium and nitrate nutrition in irrigated and rainfed environments. The study further evaluated GS and GOGAT activity and examined the expression of selected nitrogen assimilation genes in contrasting rice genotypes under hydroponic conditions. This integrated approach was used to identify genotypes with contrasting responses to nitrogen forms and water availability.

## 2. Materials and Methods

A total of 48 recombinant inbred lines (RILs), developed from a cross between Danteshwari—a high-yielding, dwarf, and drought-susceptible indica variety—and Dagaddeshi—a tall, early-maturing, drought-tolerant indica landrace— were evaluated under varying moisture regimes (irrigated and rainfed) and nitrogen forms (NH and NO ) across two cropping seasons. The treatment combinations applied across all environmental conditions are detailed in Table 1. The rainfed experimental fields were upland in topology, characterized by adequate drainage and percolation capacity (Table 1).

**Table 1.** Treatments for studying drought tolerance along with NH4+ and NO3-use efficiency in selected 48 RIL population.

| Levels of first factor | Fertilizer | Treatment |
| --- | --- | --- |
| N1 | Ammonium sulfate | $\text{NH}_4^+$-N |
| N2 | Calcium Nitrate | $\text{NO}_3^-$-N |
| N3 | Control | $\text{N}^0$-N |
| Levels of second factor | Environmental/field conditions | Soil moisture regime |
| E1 | Irrigated/Anaerobic | Non-stress |
| E2 | Rainfed/Aerobic | Stress |

The experiment followed a factorial randomized block design with two replications. Three representative samples were selected for recording the traits to obtain absolute accuracy. Agronomic, physiological, and biochemical parameters were recorded, including grain yield, plant height, leaf rolling, leaf color chart (LCC) scores, and chlorophyll content measured using a SPAD meter. Selection of superior lines was based on performance under the imposed conditions, with emphasis on traits such as SPAD readings, chlorophyll content, grain yield, and plant height.

The data obtained under different trials were tabulated and subjected to statistical analysis using Proc Means of Statistical Analysis System (SAS) v.9.1. Data were analyzed by analysis of variance, and the F-test was used to determine the treatment significance. The mean data of each replication was used for analysis of variance using factorial design due to the presence of multiple variables in the experiments.

The activities of key nitrogen assimilation enzymes—glutamine synthetase (GS) and glutamate synthase (GOGAT)—were assayed in freshly harvested leaves from 48 selected rice genotypes grown under field conditions. Enzyme activity measurements were conducted at the maximum tillering to flowering stage. Plants were treated with calcium nitrate and ammonium sulfate in the recommended agronomic ratio, alongside untreated controls. Enzyme extractions were performed following the protocol described by (Sadasivam 1996). Each enzymatic assay was conducted in duplicate, and the results were expressed as relative specific activity (%) with associated standard errors. Specific enzyme activity was defined as the amount of product formed (µmol) per milligram of protein.

### Expression analysis of genes involved in nitrogen metabolism

The RT-PCR-based expression analysis of assimilatory genes Glutamine synthetase (GS) and Glutamate synthase (GOGAT) were carried out on four selected rice genotypes at tillering stage. Details of the selected genotypes and the selected primers for the RT-PCR study are listed in Table 2 and 3. The entire experimental setup was established under hydroponic conditions where nitrogen treatments were given in the form of ammonium sulfate, calcium nitrate, full-strength ammonium nitrate and half-strength ammonium nitrate treatments. For the targeted gene expression analysis of selected nitrogen metabolism genes, both root and leaf materials were harvested from two biological replicates. Total RNA was extracted using Pure LinkTM RNA mini-Kit (Invitrogen) and was used for cDNA synthesis using Thermo Scientific VersoTM cDNA Synthesis kit. The cDNA samples were used as templates to quantify target gene expression levels in individual genotypes under a particular treatment. The qPCR was performed using gene-specific primers with actin as an internal control. qPCR was accomplished on Mx3000P qPCR system (Stratagene, USA).

**Table 2.** Lines selected for expression analysis studies.

| Genotype No. | Sample ID | Leaf colour |
| --- | --- | --- |
| 1 | Danteshwari | Green |
| 2 | Dagaddeshi | Light green |
| 3 | Swarna | Dark green |
| 4 | Indira Sugandhit Dhan 1 (ISD-1) | Yellow |

**Table 3.**
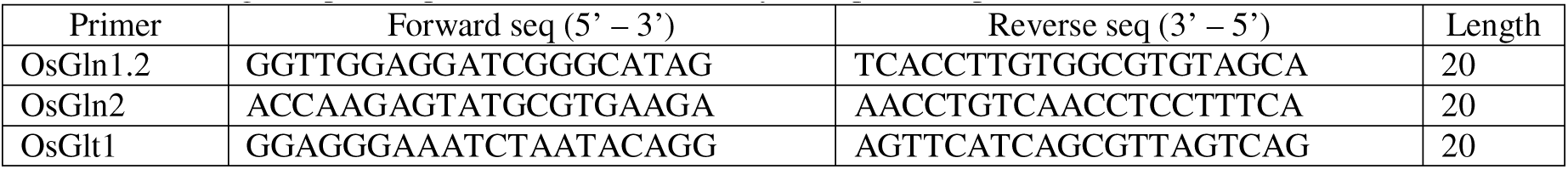
List of gene specific primers used for the study of expression profile.

## 3. Results

### 3.1. Phenotyping

Evaluation of morphological traits in the rice accessions revealed a wide range of variation among the genotypes for all the measured yield and yield-related parameters. The data recorded for all the experiments were subjected to a three-factor analysis of variance, and the mean sum of squares due to various sources of variation for the investigated traits are summarized in Tables 4 and 5. Analysis of Variance (ANOVA) showed individual sources of variation, such as genotype (G), nitrogen (N), and environment (E), are significant. Analysis of variance revealed significant effects of genotype, nitrogen treatment, and environment on several traits, with significant interactions among these factors for many traits.

**Table 4.** Mean sum of squares (MSS) of variance components and their interactions of selected 48 RILs along with parents under NH_4_^+^ and NO_3_^-^ forms of nitrogen and variable environments for yield and yield related traits during second wet season .

| Traits | Wet season II |  |  |  |  |  |  |
| --- | --- | --- | --- | --- | --- | --- | --- |
|  | G | N | E | E X G | N X G | EX N | E X N X G |
|  | TMSS, DF=47 | TMSS, DF=2 | TMSS, DF=1 | TMSS, DF=47 | TMSS, DF=94 | TMSS, DF=2 | TMSS, DF=94 |
| GY | 299487.24*** | 657839.95*** | 1831398.33*** | 146482.65*** | 22251.02** | 753308.7*** | 1,358,658.54* |
| BY | 1216262.23*** | 17755402.45*** | 3436389.06*** | 366482.74*** | 102287.65** | 2487375.95*** | 7,265,406.79* |
| HI | 744.21*** | 1804.79*** | 1198.52*** | 517.45*** | 45.389 | 956.30*** | 3875.169* |
| DTF | 413** | 4,619** | 995** | 358** | 445** | 1,234** | 35620** |
| PH | 7135.61*** | 2246.59*** | 40614.80*** | 282.41*** | NS | 1799.24*** | 45.521* |
| PL | 41.91*** | 6.799** | 1091.9*** | 10.08*** | 3.65*** | 101.04*** | 3.149** |
| FLA | 456.50*** | 2143.21*** | 3101.23*** | 106.68** | 72.43* | 536.23*** | 80.81* |
| FLL | 260.31*** | 569.87*** | 8290.98*** | 48.38*** | 16.26* | 43.27** | 17.22*** |
| FLW | 0.26*** | 2.34*** | 1.19*** | NS | NS | 1.44*** | 0.11** |
| PLL | 816.64*** | 942.94*** | 942.94*** | 160.37*** | 47.87*** | 85*** | 31.54** |
| PLW | 0.23*** | 1.69*** | 0.087* | 0.073*** | 0.056*** | 1.59*** | 1.56*** |
| SL | 5.75*** | 0.584*** | 0.963*** | 0.31*** | 0.377*** | 0.016* | 0.182*** |
| SB | 0.643*** | 2.546*** | 0.202*** | 0.131*** | 0.089*** | 0.258*** | 0.053*** |
| SLB | 1.708*** | 2.342*** | 0.715*** | 0.25*** | 0.206*** | 0.552*** | 0.12*** |
| 100 SW | 0.773*** | 0.065*** | 0.525*** | 0.06*** | 0.05*** | 0.029*** | 0.045*** |
| SF | 787.50*** | 266.42*** | 6619.91*** | 441.26*** | 89.88*** | 850.78*** | 118.03*** |
| SS | 787.38*** | 263.39*** | 6614.1*** | 441.38*** | 89.94*** | 853.71*** | 117.97*** |
| Total Chlorophyll | 25.03*** | 299.31*** | 2.38* | 15.88*** | 11.049*** | 1006.31*** | 13.078*** |

\*, \*\* and \*\*\* significance at .05 .01 and .001 levels respectively, GY=grain yield, BG= biological yield, HI= harvest index, PH= plant height, PL=panicle length, FLL=flag leaf length, FLW=flag leaf width, FLA=flag leaf area, PLL= penultimate leaf length, PLW= penultimate leaf width, SL= seed length, SB= seed breadth, SLB= seed length and breadth ratio, SF= spikelet fertility, SS= spikelet sterility.

**Table 5.**
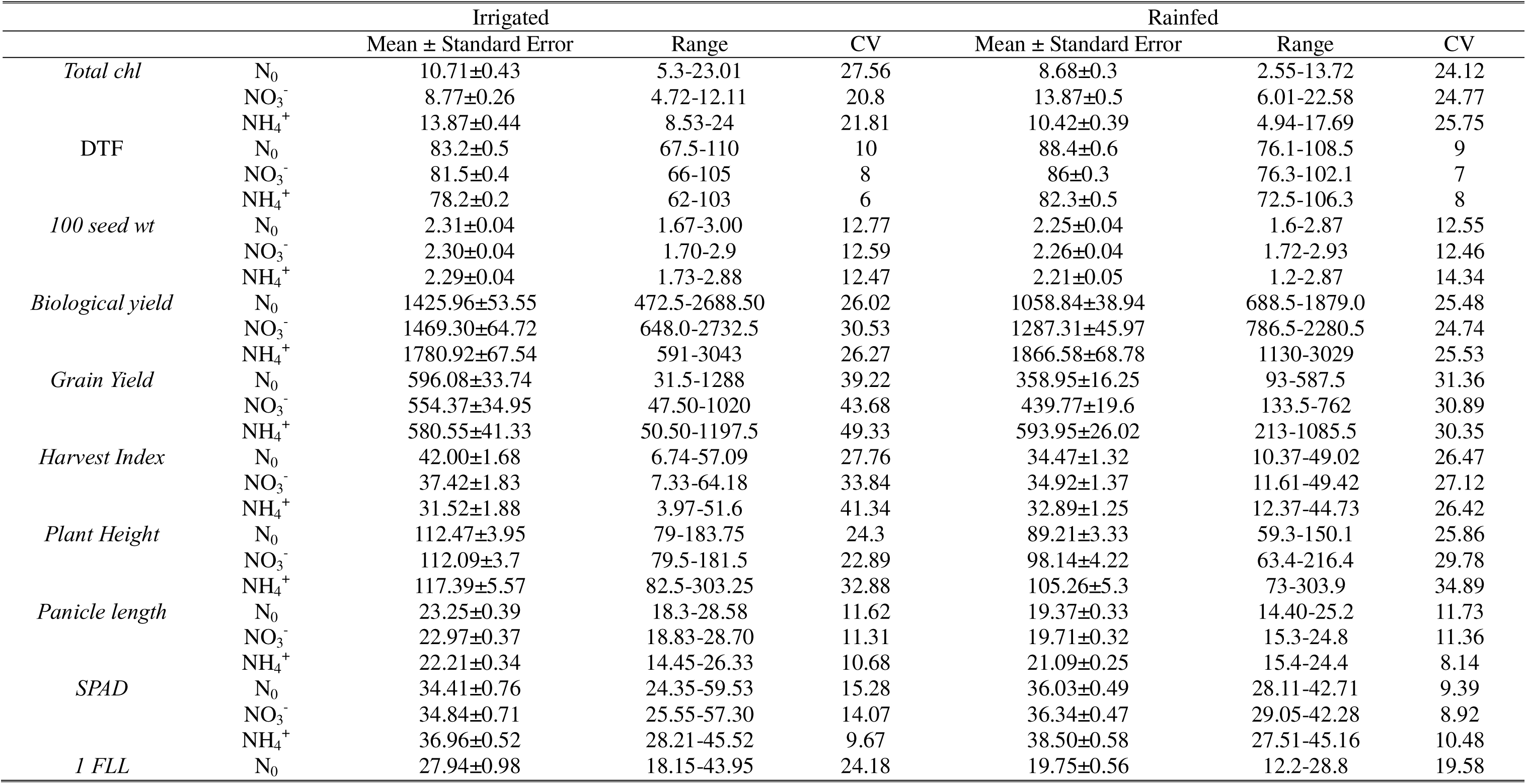
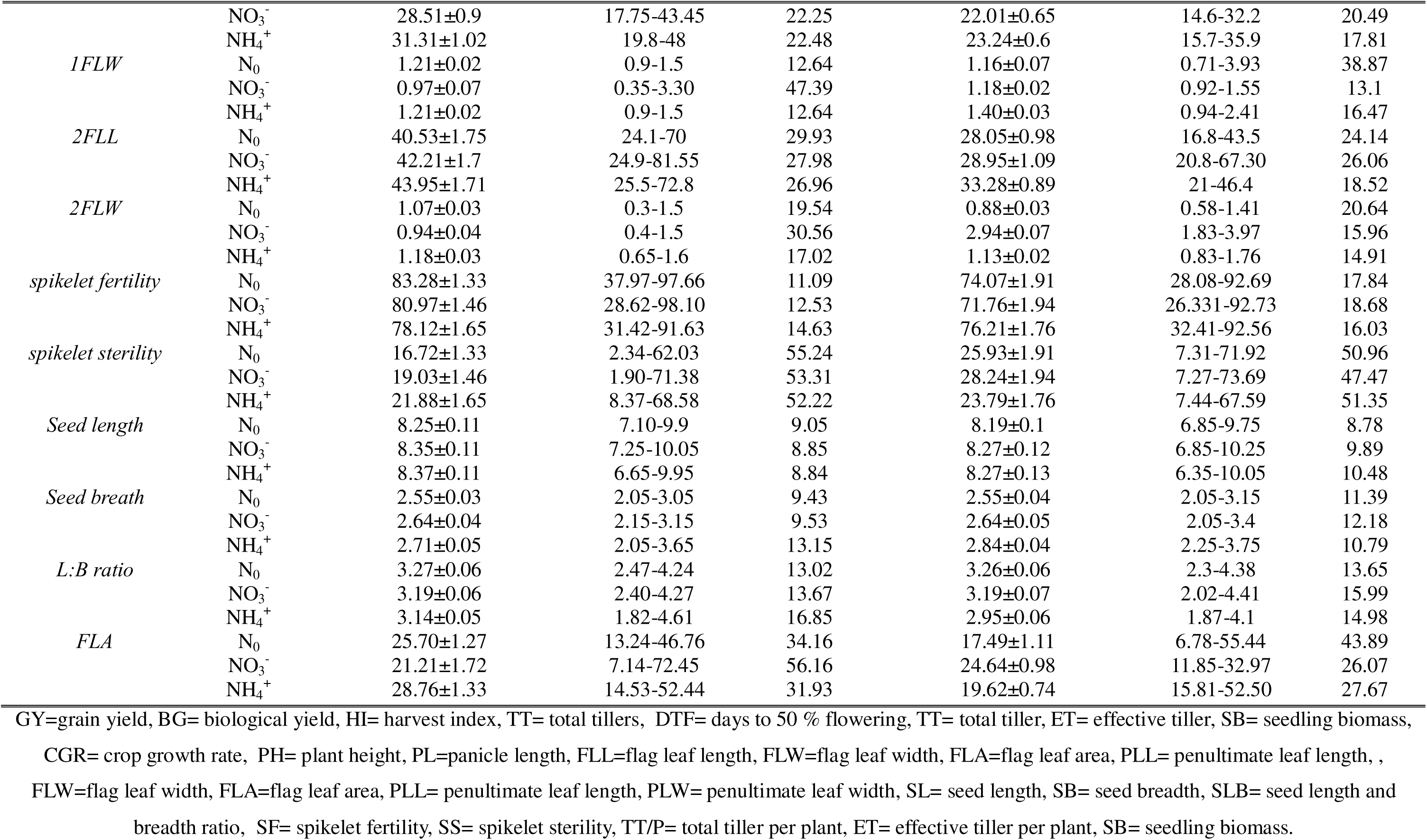
Mean phenotypic performance, range, Coefficient of variance (CV %) of investigated yield and yield-related traits of 48 selected RILs under differential N regimes and environments during the second wet season

### 3.2. Agronomic traits under different nitrogen forms

There was a significant reduction in the grain yield under rainfed conditions compared to the irrigated environment. In 48 RILs, the coefficient of variation for GY under different water regimes ranged from 30.35 to 49.33% and higher grain yields were observed with ammoniacal nitrogen than nitrate nitrogen under rainfed conditions, but under irrigated conditions, the control mean was slightly higher than ammonium. Biological yield had the highest mean under ammonium treatment in both irrigated and rainfed environments. Harvest Index (HI) was higher in control (no nitrogen) treatment under irrigated environments. Mean HI was higher in nitrate form in the 48 RILs, indicating a high variation in this trait. Earlier flowering was observed under ammoniacal treatment, both in rainfed and irrigated conditions.

Soil-Plant Analysis Development (SPAD) and leaf chlorophyll content can give information on the nitrogen status of the plant. In the present study, we observed significant differences in leaf chlorophyll content and SPAD meter values under different forms of nitrogen at flourishing tillering stage (Table **6**). Chlorophyll and SPAD values were subjected to ANOVA revealed that the genotype and nitrogen forms have individual effect, and their interactions are also significant. The effect of the environment on SPAD was non-significant in season 1 but significant in season 2. Total chlorophyll showed a treatment and season-dependent response. Under irrigated conditions, ammonium produced the highest mean chlorophyll content in both seasons. Under rainfed conditions, ammonium produced the highest mean in the first season, whereas nitrate produced the highest mean in the second season. SPAD values were consistently highest under ammonium treatment. The corresponding R2 values with NH4+, NO3-and treatment under irrigated trials were 0.7, 0.670, and 0.657 respectively and in rainfed conditions, the values were 0.633, 0.758 and 0.587 during the first wet season. During the second season, the R2 values for the NH4+, NO3-, and control treatments in the irrigated trial were 0.786, 0.729, and 0.812, respectively, and under rainfed conditions, the values were 0.672, 0.797, and 0.707, respectively.

**Table 6.**
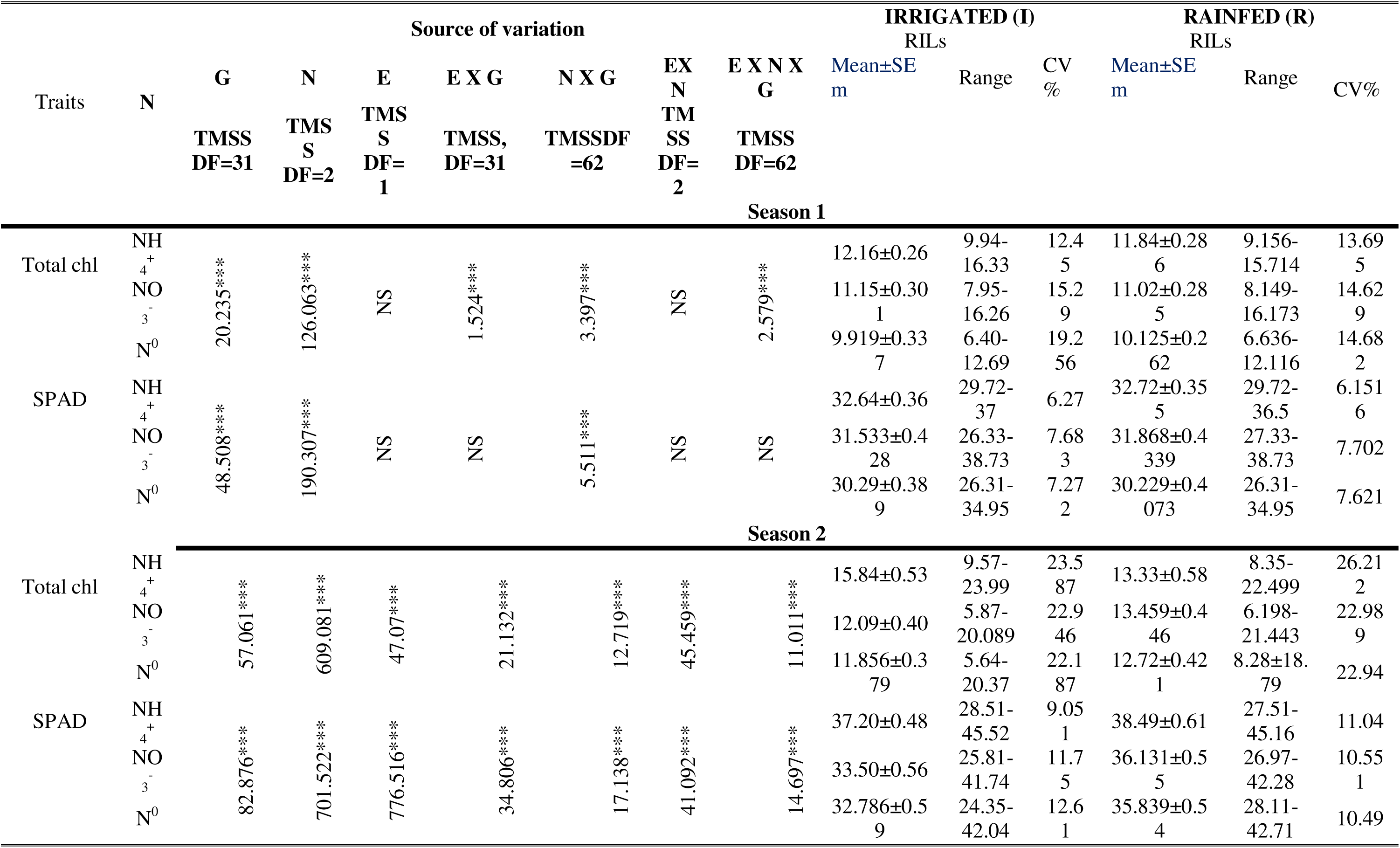
Mean phenotypic performance, range, standard deviation (SD), Coefficient of variance (CV %) of investigated SPAD and chlorophyll parameters of 48 selected RILs and their parents under differential N regimes and environments during the wet season, first and second season, respectively.

### 3.3. Comparison of grain yield under water regimes with different forms of nitrogen

Across the 48 rice genotypes, grain-yield responses to nitrogen varied with both genotype and water regime. Under irrigated conditions, the mean grain yield was slightly higher with ammonium than nitrate (580.55 vs 554.37 g m^-2^), whereas under rainfed conditions, the difference was more pronounced (593.95 vs 439.77 g m^-2^). Ammonium produced higher grain yield than nitrate in 23 of the 48 genotypes under irrigation and in 44 genotypes under rainfed conditions (Table 7). Thus, the yield advantage associated with ammonium was greater under rainfed conditions, although the response was genotype dependent.

**Table 7.**
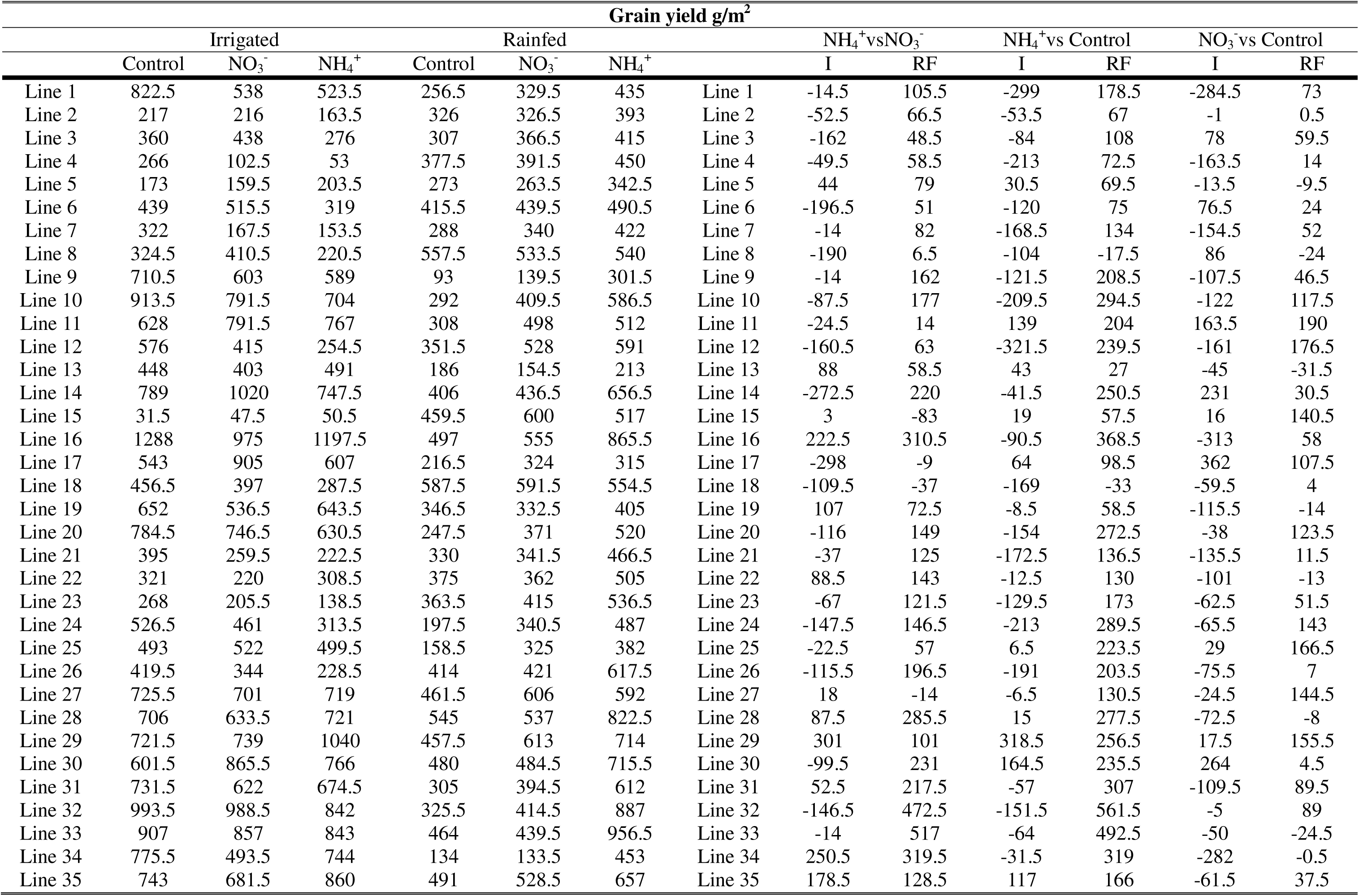
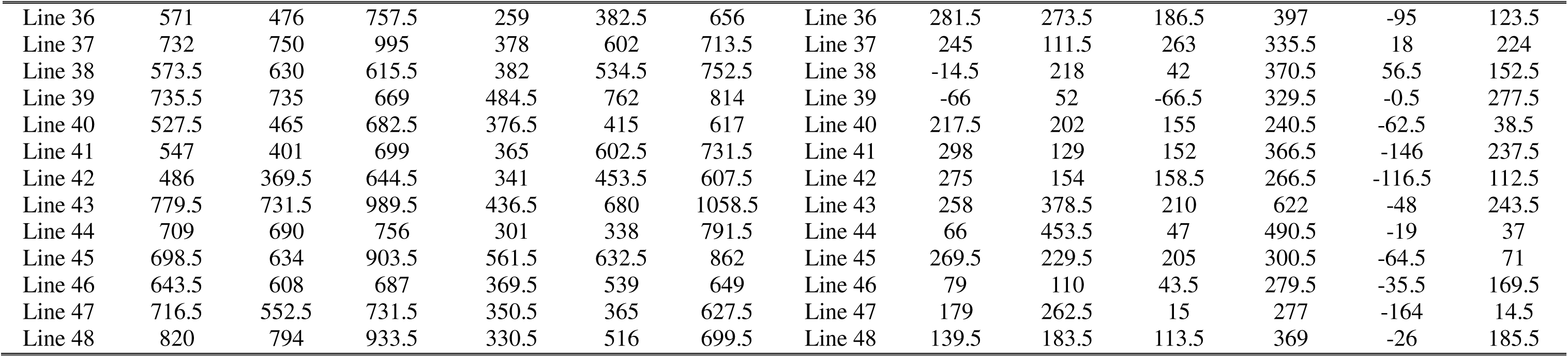
Comparison of yield of selected 48 rice genotypes under different N regimes.

| Grain yield g/m<sup>2</sup> |  |  |  |  |  |  |  |  |  |  |  |  |  |
| --- | --- | --- | --- | --- | --- | --- | --- | --- | --- | --- | --- | --- | --- |
|  | Irrigated |  |  | Rainfed |  |  |  | NH<sub>4</sub><sup>+</sup> vs NO<sub>3</sub><sup>-</sup> |  | NH<sub>4</sub><sup>+</sup> vs Control |  | NO<sub>3</sub><sup>-</sup> vs Control |  |
|  | Control | NO<sub>3</sub><sup>-</sup> | NH<sub>4</sub><sup>+</sup> | Control | NO<sub>3</sub><sup>-</sup> | NH<sub>4</sub><sup>+</sup> |  | I | RF | I | RF | I | RF |
| Line 1 | 822.5 | 538 | 523.5 | 256.5 | 329.5 | 435 | Line 1 | -14.5 | 105.5 | -299 | 178.5 | -284.5 | 73 |
| Line 2 | 217 | 216 | 163.5 | 326 | 326.5 | 393 | Line 2 | -52.5 | 66.5 | -53.5 | 67 | -1 | 0.5 |
| Line 3 | 360 | 438 | 276 | 307 | 366.5 | 415 | Line 3 | -162 | 48.5 | -84 | 108 | 78 | 59.5 |
| Line 4 | 266 | 102.5 | 53 | 377.5 | 391.5 | 450 | Line 4 | -49.5 | 58.5 | -213 | 72.5 | -163.5 | 14 |
| Line 5 | 173 | 159.5 | 203.5 | 273 | 263.5 | 342.5 | Line 5 | 44 | 79 | 30.5 | 69.5 | -13.5 | -9.5 |
| Line 6 | 439 | 515.5 | 319 | 415.5 | 439.5 | 490.5 | Line 6 | -196.5 | 51 | -120 | 75 | 76.5 | 24 |
| Line 7 | 322 | 167.5 | 153.5 | 288 | 340 | 422 | Line 7 | -14 | 82 | -168.5 | 134 | -154.5 | 52 |
| Line 8 | 324.5 | 410.5 | 220.5 | 557.5 | 533.5 | 540 | Line 8 | -190 | 6.5 | -104 | -17.5 | 86 | -24 |
| Line 9 | 710.5 | 603 | 589 | 93 | 139.5 | 301.5 | Line 9 | -14 | 162 | -121.5 | 208.5 | -107.5 | 46.5 |
| Line 10 | 913.5 | 791.5 | 704 | 292 | 409.5 | 586.5 | Line 10 | -87.5 | 177 | -209.5 | 294.5 | -122 | 117.5 |
| Line 11 | 628 | 791.5 | 767 | 308 | 498 | 512 | Line 11 | -24.5 | 14 | 139 | 204 | 163.5 | 190 |
| Line 12 | 576 | 415 | 254.5 | 351.5 | 528 | 591 | Line 12 | -160.5 | 63 | -321.5 | 239.5 | -161 | 176.5 |
| Line 13 | 448 | 403 | 491 | 186 | 154.5 | 213 | Line 13 | 88 | 58.5 | 43 | 27 | -45 | -31.5 |
| Line 14 | 789 | 1020 | 747.5 | 406 | 436.5 | 656.5 | Line 14 | -272.5 | 220 | -41.5 | 250.5 | 231 | 30.5 |
| Line 15 | 31.5 | 47.5 | 50.5 | 459.5 | 600 | 517 | Line 15 | 3 | -83 | 19 | 57.5 | 16 | 140.5 |
| Line 16 | 1288 | 975 | 1197.5 | 497 | 555 | 865.5 | Line 16 | 222.5 | 310.5 | -90.5 | 368.5 | -313 | 58 |
| Line 17 | 543 | 905 | 607 | 216.5 | 324 | 315 | Line 17 | -298 | -9 | 64 | 98.5 | 362 | 107.5 |
| Line 18 | 456.5 | 397 | 287.5 | 587.5 | 591.5 | 554.5 | Line 18 | -109.5 | -37 | -169 | -33 | -59.5 | 4 |
| Line 19 | 652 | 536.5 | 643.5 | 346.5 | 332.5 | 405 | Line 19 | 107 | 72.5 | -8.5 | 58.5 | -115.5 | -14 |
| Line 20 | 784.5 | 746.5 | 630.5 | 247.5 | 371 | 520 | Line 20 | -116 | 149 | -154 | 272.5 | -38 | 123.5 |
| Line 21 | 395 | 259.5 | 222.5 | 330 | 341.5 | 466.5 | Line 21 | -37 | 125 | -172.5 | 136.5 | -135.5 | 11.5 |
| Line 22 | 321 | 220 | 308.5 | 375 | 362 | 505 | Line 22 | 88.5 | 143 | -12.5 | 130 | -101 | -13 |
| Line 23 | 268 | 205.5 | 138.5 | 363.5 | 415 | 536.5 | Line 23 | -67 | 121.5 | -129.5 | 173 | -62.5 | 51.5 |
| Line 24 | 526.5 | 461 | 313.5 | 197.5 | 340.5 | 487 | Line 24 | -147.5 | 146.5 | -213 | 289.5 | -65.5 | 143 |
| Line 25 | 493 | 522 | 499.5 | 158.5 | 325 | 382 | Line 25 | -22.5 | 57 | 6.5 | 223.5 | 29 | 166.5 |
| Line 26 | 419.5 | 344 | 228.5 | 414 | 421 | 617.5 | Line 26 | -115.5 | 196.5 | -191 | 203.5 | -75.5 | 7 |
| Line 27 | 725.5 | 701 | 719 | 461.5 | 606 | 592 | Line 27 | 18 | -14 | -6.5 | 130.5 | -24.5 | 144.5 |
| Line 28 | 706 | 633.5 | 721 | 545 | 537 | 822.5 | Line 28 | 87.5 | 285.5 | 15 | 277.5 | -72.5 | -8 |
| Line 29 | 721.5 | 739 | 1040 | 457.5 | 613 | 714 | Line 29 | 301 | 101 | 318.5 | 256.5 | 17.5 | 155.5 |
| Line 30 | 601.5 | 865.5 | 766 | 480 | 484.5 | 715.5 | Line 30 | -99.5 | 231 | 164.5 | 235.5 | 264 | 4.5 |
| Line 31 | 731.5 | 622 | 674.5 | 305 | 394.5 | 612 | Line 31 | 52.5 | 217.5 | -57 | 307 | -109.5 | 89.5 |
| Line 32 | 993.5 | 988.5 | 842 | 325.5 | 414.5 | 887 | Line 32 | -146.5 | 472.5 | -151.5 | 561.5 | -5 | 89 |
| Line 33 | 907 | 857 | 843 | 464 | 439.5 | 956.5 | Line 33 | -14 | 517 | -64 | 492.5 | -50 | -24.5 |
| Line 34 | 775.5 | 493.5 | 744 | 134 | 133.5 | 453 | Line 34 | 250.5 | 319.5 | -31.5 | 319 | -282 | -0.5 |
| Line 35 | 743 | 681.5 | 860 | 491 | 528.5 | 657 | Line 35 | 178.5 | 128.5 | 117 | 166 | -61.5 | 37.5 |
| Line 36 | 571 | 476 | 757.5 | 259 | 382.5 | 656 | Line 36 | 281.5 | 273.5 | 186.5 | 397 | -95 | 123.5 |
| Line 37 | 732 | 750 | 995 | 378 | 602 | 713.5 | Line 37 | 245 | 111.5 | 263 | 335.5 | 18 | 224 |
| Line 38 | 573.5 | 630 | 615.5 | 382 | 534.5 | 752.5 | Line 38 | -14.5 | 218 | 42 | 370.5 | 56.5 | 152.5 |
| Line 39 | 735.5 | 735 | 669 | 484.5 | 762 | 814 | Line 39 | -66 | 52 | -66.5 | 329.5 | -0.5 | 277.5 |
| Line 40 | 527.5 | 465 | 682.5 | 376.5 | 415 | 617 | Line 40 | 217.5 | 202 | 155 | 240.5 | -62.5 | 38.5 |
| Line 41 | 547 | 401 | 699 | 365 | 602.5 | 731.5 | Line 41 | 298 | 129 | 152 | 366.5 | -146 | 237.5 |
| Line 42 | 486 | 369.5 | 644.5 | 341 | 453.5 | 607.5 | Line 42 | 275 | 154 | 158.5 | 266.5 | -116.5 | 112.5 |
| Line 43 | 779.5 | 731.5 | 989.5 | 436.5 | 680 | 1058.5 | Line 43 | 258 | 378.5 | 210 | 622 | -48 | 243.5 |
| Line 44 | 709 | 690 | 756 | 301 | 338 | 791.5 | Line 44 | 66 | 453.5 | 47 | 490.5 | -19 | 37 |
| Line 45 | 698.5 | 634 | 903.5 | 561.5 | 632.5 | 862 | Line 45 | 269.5 | 229.5 | 205 | 300.5 | -64.5 | 71 |
| Line 46 | 643.5 | 608 | 687 | 369.5 | 539 | 649 | Line 46 | 79 | 110 | 43.5 | 279.5 | -35.5 | 169.5 |
| Line 47 | 716.5 | 552.5 | 731.5 | 350.5 | 365 | 627.5 | Line 47 | 179 | 262.5 | 15 | 277 | -164 | 14.5 |
| Line 48 | 820 | 794 | 933.5 | 330.5 | 516 | 699.5 | Line 48 | 139.5 | 183.5 | 113.5 | 369 | -26 | 185.5 |

### 3.4. Correlation between grain yield and yield contributing traits of rice genotypes under variable nutrient levels

A total of 16 pair wise combinations under six conditions (formed from combination of two environmental conditions and three treatments) of grain yield were analyzed. Correlation between these traits was presented in Table 8. Highly significant and positive correlation between biological yield and grain yield observed under irrigated with ammoniacal, nitrate and control treatment were 0.928, 0.930 and 0.884 respectively and 0.917, 0.690 and 0.763 under rainfed conditions. Plant height showed a significant and positive correlation with grain yield under control (0.359*) and nitrate (0.381*) conditions in irrigated conditions and in rainfed control (r=0.430**). Grain yield showed significant positive correlations with selected flag leaf traits, including second flag leaf length under irrigated nitrate nutrition and first and second flag leaf width under rainfed ammonium nutrition. Seed length and seed length: breadth ratio showed negative correlation with grain yield under ammoniacal conditions in both environments.

**Table 8.**
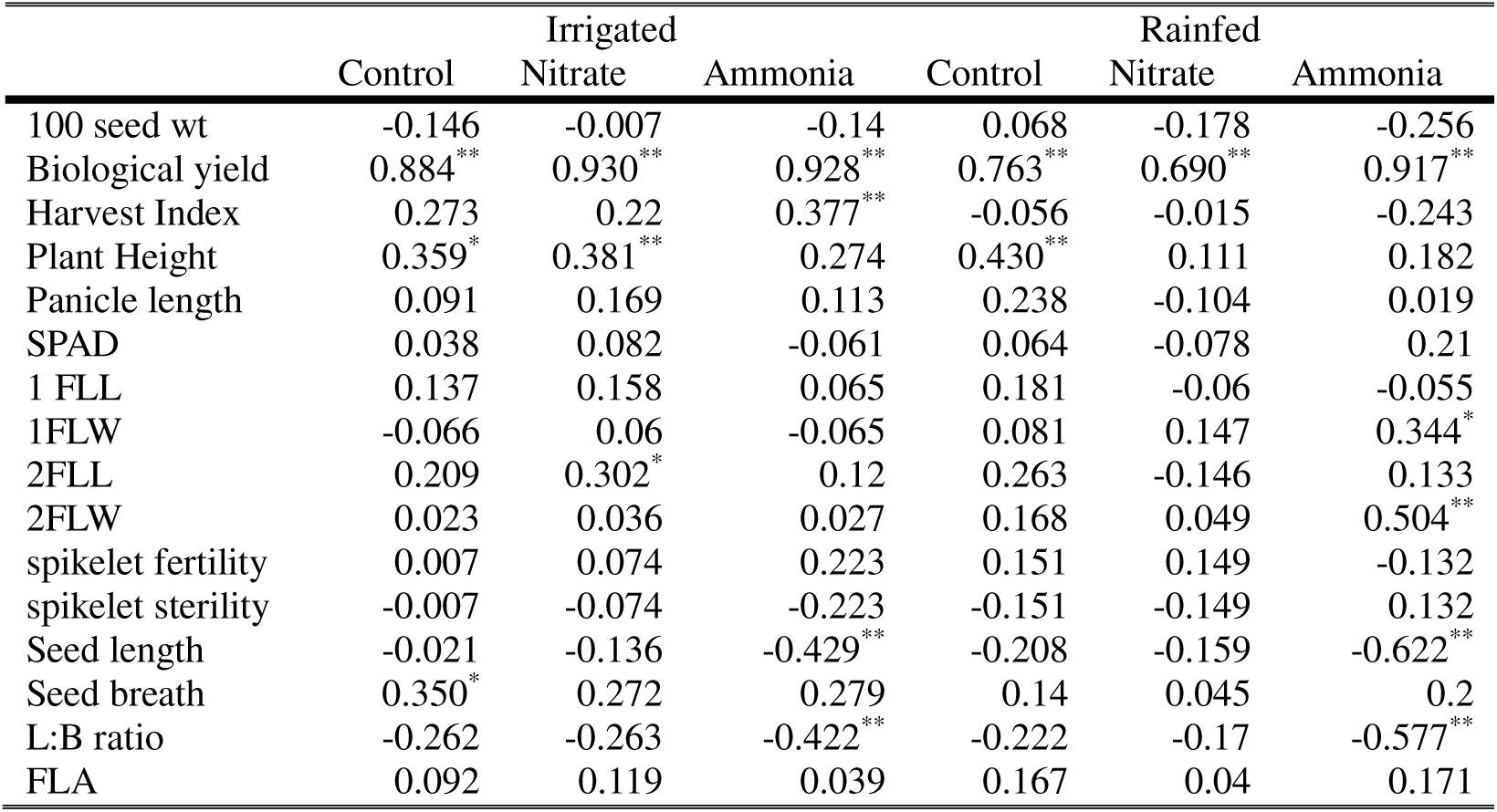
Correlation between grain yield and yield contributing traits under differential N regimes under irrigated and rainfed conditions.

### 3.5. Enzymatic activities and their relationship grain yield

The activity of major enzymes involved in nitrogen assimilation; Glutamine synthetase (GS), Glutamate synthetase (GOGAT) were measured in the selected 48 genotypes rice genotypes at maximum tillering stage under rainfed condition and were analyzed by standard analysis of variance techniques to assess the significance of treatment means. Enzyme activity data generated were represented as mean values ±SE_m,_ range (minimum-maximum) and CV value. The genotypes exhibited variance for most of the phenotypic traits studied. The results of the present study showed that nitrogen forms greatly influence enzymatic activity, and the results are depicted in Table 9. ANOVA showed significant variation due to genotypes, nitrogen forms, and their combination, which showed that ammoniacal form of nitrogen control GS activity whereas GOGAT activity was higher under nitrate treatment. The highest mean GS activity was recorded under ammoniacal form followed by nitrate and control treatment. The coefficient of variation ranges between 15.58 to 22.18% for GS activity. Highest GS activity was observed in DxD 124-21 (line #24) in NH_4_^+^ treatment, 12.1(487)-237 (line #40) in NO_3_^-^ and MTU1010 (line #10) in control treatment. The high yielding varieties like Swarna (line #4), IBD (line #19) were found to have high GS activity under both ammonium and nitrate treatments (Fig. 1).

**Fig 1.**
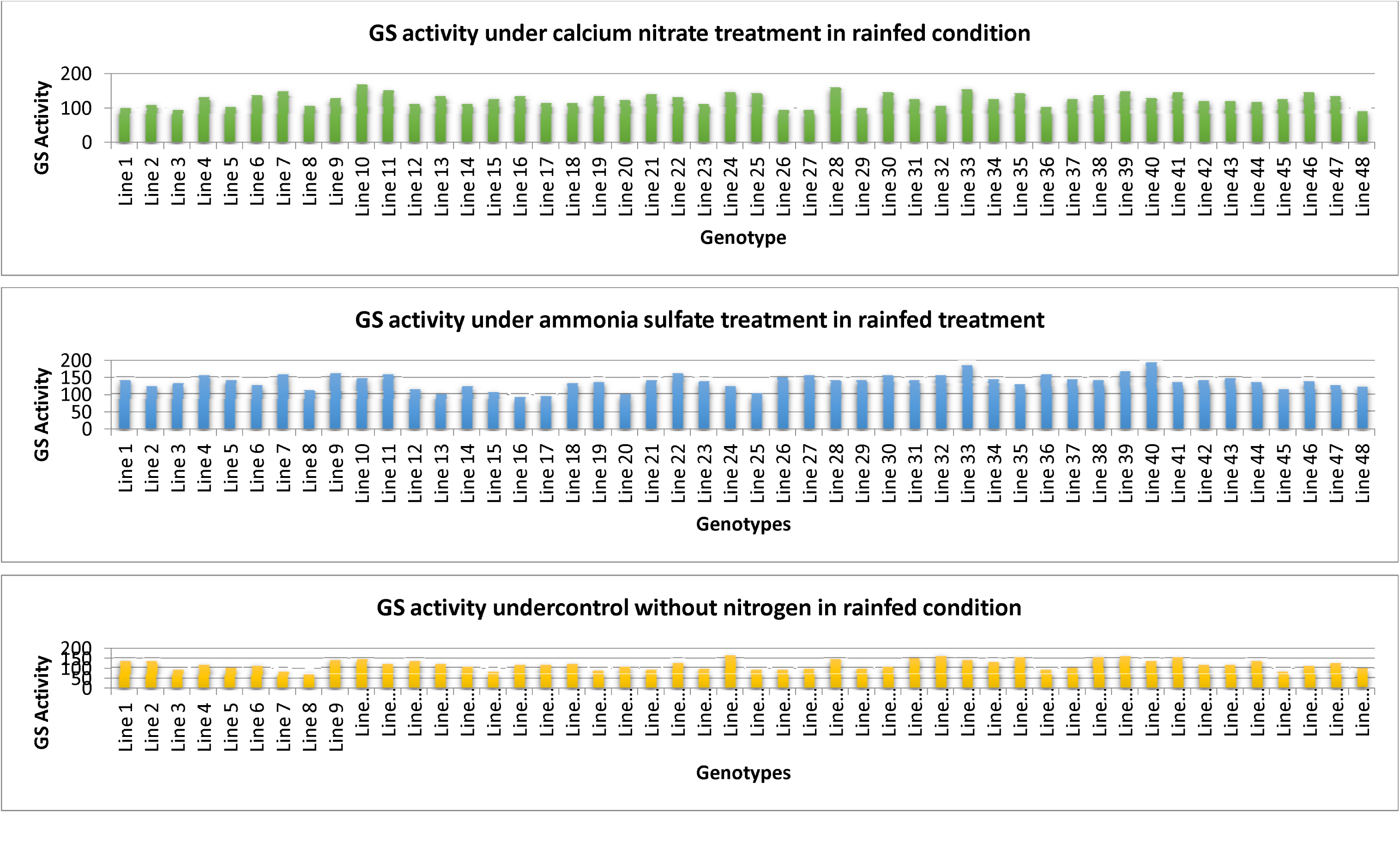
Glutamine Synthetase activity in rice genotypes grown under rainfed conditions.

**Table 9.**
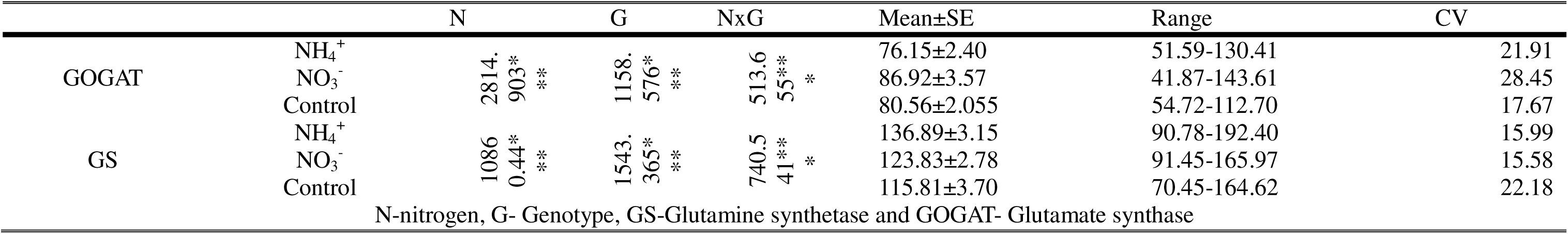
ANOVA and mean phenotypic performance, range, Coefficient of variance (CV %) of GS and GOGAT activity in 48 selected RILs with parents under differential N regimes during the second wet season

The GOGAT activity followed a different pattern from that of GS (Fig. 2). The highest mean GOGAT activity was recorded in leaf tissues under NO_3_^-^ treatment over NH_4_^+^ and N_0_ (control) treatment. The coefficient of variation ranged between 17.67 to 28.45% for GOGAT activity. The highest GOGAT activity was observed in RKVY-104 (line #18) in NO_3_^-^ treatment, DxD 124-75 (line #5) and GP 145-42 (line #14) in control treatment. Grain yield showed positive but nonsignificant correlations with GS activity under ammonium and nitrate treatments, whereas the correlation was negative and nonsignificant under the control treatment. GOGAT activity showed negative and nonsignificant correlations with grain yield under all nitrogen treatments (Table 10).

**Fig. 2.**
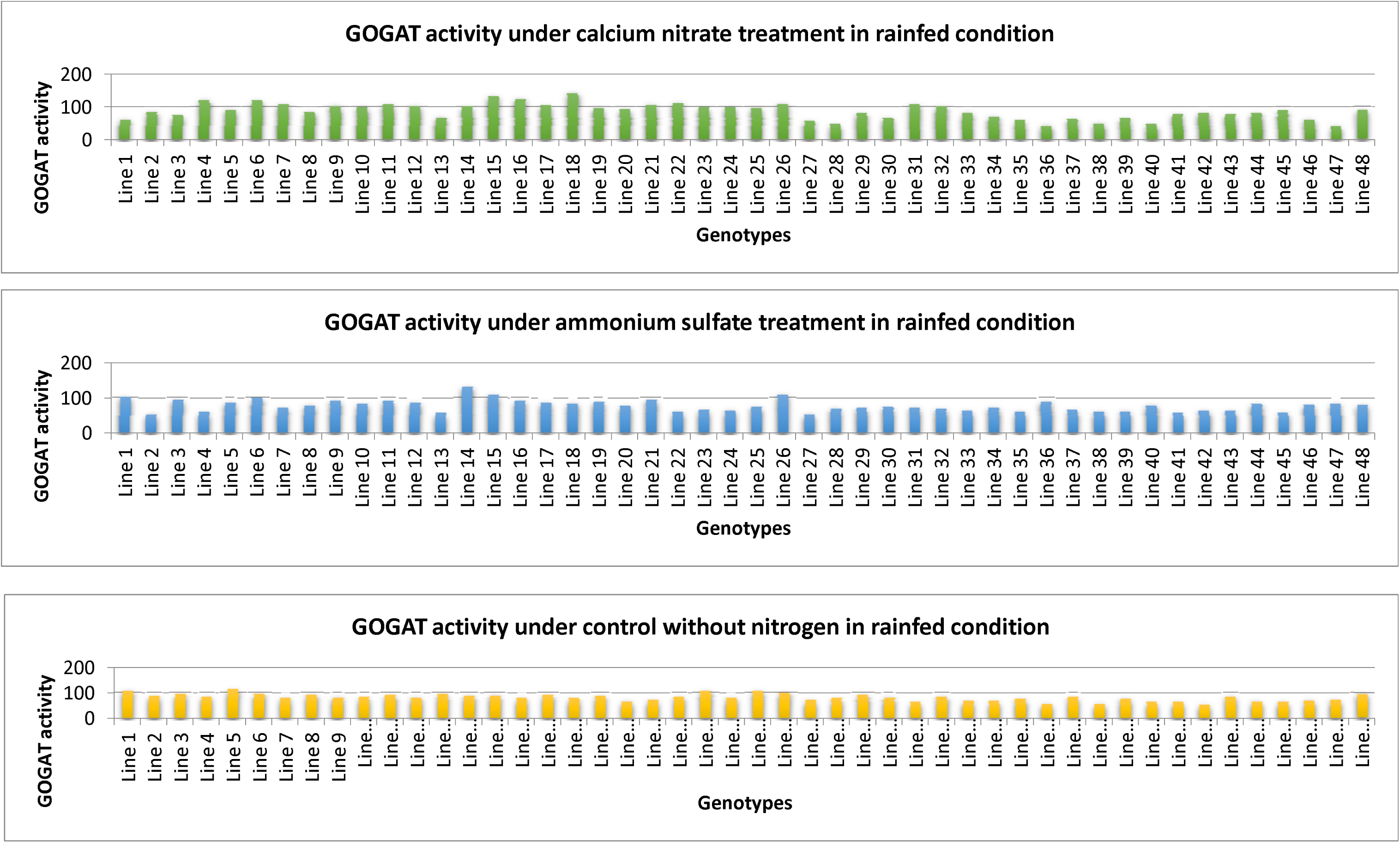
Glutamate Synthase activity in rice genotypes grown under rainfed condition.

**Table 10.** Correlation studies between grain yield and enzyme activity under different forms of nitrogen treatment in second wet season GS activity GOGAT activity.

|  | GS activity |  |  | GOGAT activity |  |  |
| --- | --- | --- | --- | --- | --- | --- |
| | $\text{NH}_4^+$ | $\text{NO}_3^-$ | $\text{N}_0$ | $\text{NH}_4^+$ | $\text{NO}_3^-$ | $\text{N}_0$ |
| GY | 0.252<sup>NS</sup> | 0.45<sup>NS</sup> | -0.135<sup>NS</sup> | -0.209<sup>NS</sup> | -0.074<sup>NS</sup> | -0.141<sup>NS</sup> |

### 3.6. Gene expression analysis

Expression of nitrogen metabolism genes varied with genotype, tissue and nitrogen treatment.

#### GS gene family

Glutamine synthetase (GS) is a major enzyme in the nitrogen assimilation pathway. GS_1_ and GS_2_ are two major isoforms of GS.

### Expression of OsGln1.2 gene

Expression of OsGln1.2 was evaluated in root and leaf tissues under ammonium sulfate, ammonium nitrate, half strength mixed nitrogen, and calcium nitrate treatments. In leaf tissues, OsGln1.2 expression was generally higher under ammonium sulfate and full-strength mixed nitrogen than under calcium nitrate and half treatment.

In root tissues, higher expression of this gene was observed under ammonium sulfate and full-strength mixed nitrogen. Expression was lower under **CN** and half treatment in several genotypes. Dagaddeshi generally showed weak expression varied according to tissue and nitrogen source (Fig. 3).

**Fig 3.**
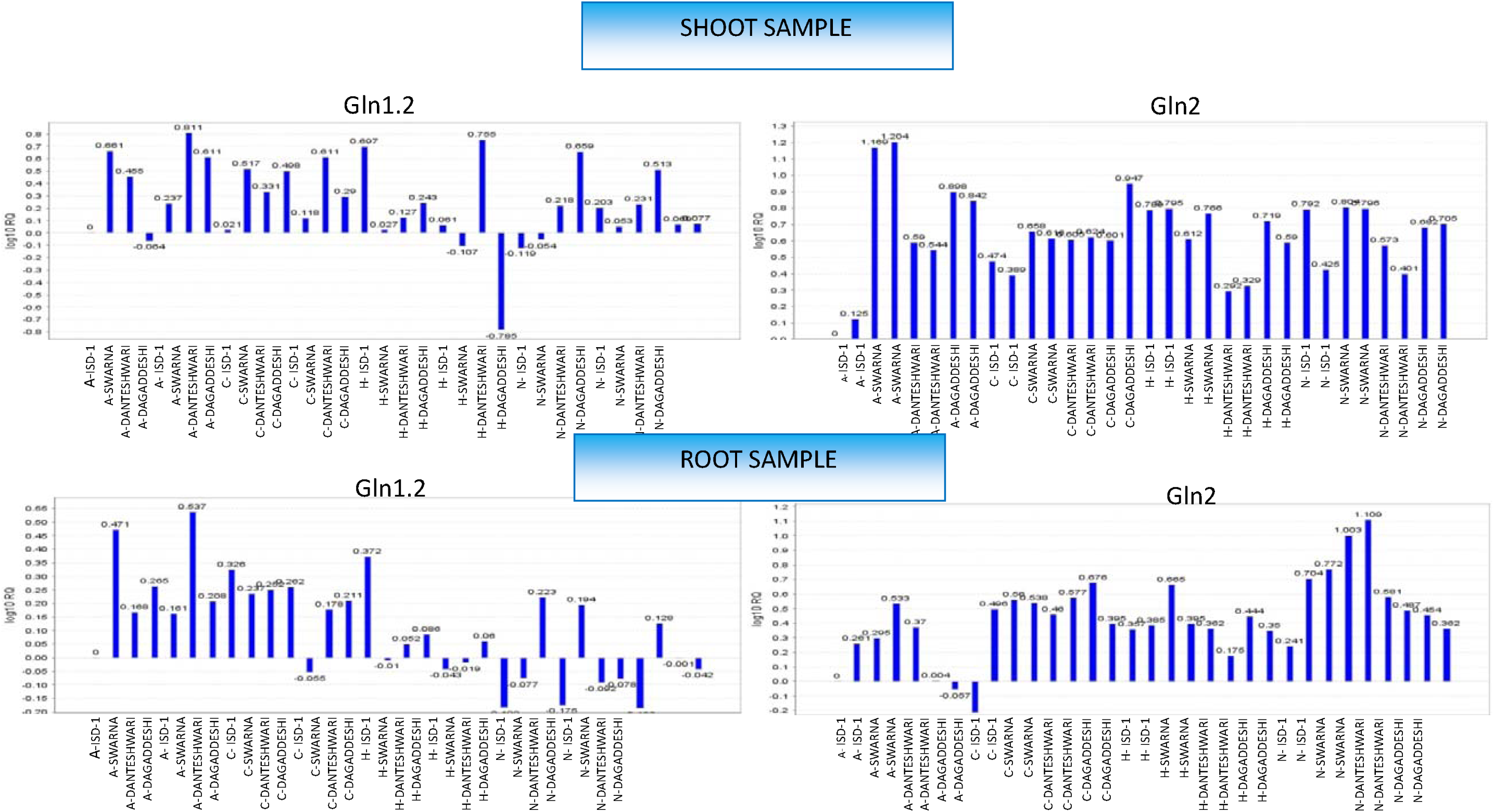
Expression profile of glutamine synthetase gene (Gln1.2 and Gln2) in shoot and root under ammonium sulfate (A), AN [Ammonium Nitrate (C)], Half [half conc. of Ammonium Nitrate (H)] treatment and calcium nitrate (N) treatment

Among the selected genotypes, Swarna and Danteshwari showed relatively high OsGln1.2 expression under ammonium sulfate and mixed nitrogen treatments. ISD-1 and Dagaddeshi showed treatment specific differences in expression between root and leaf tissues (Fig. 4).

**Fig 4.**
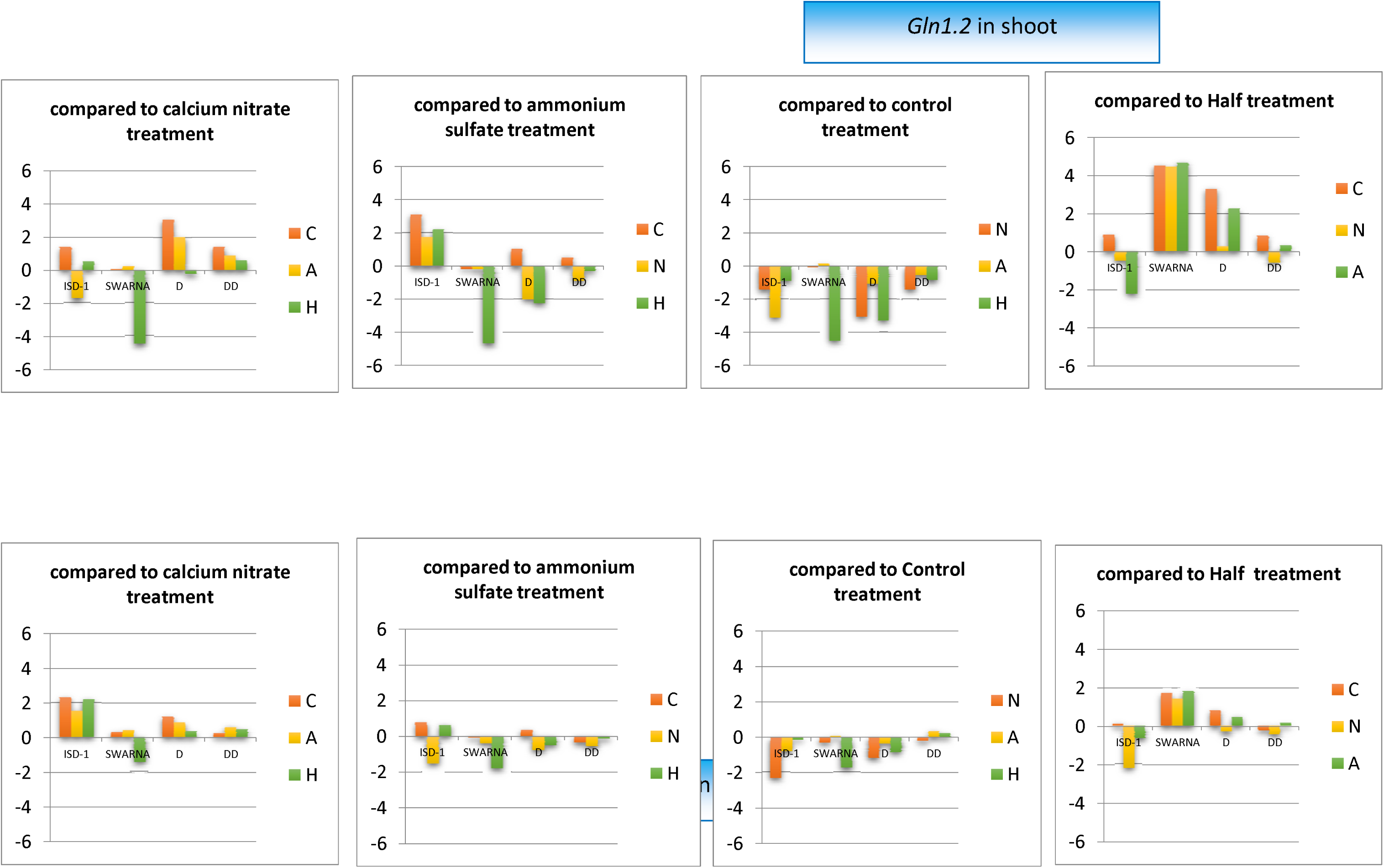
Comparison of treatments for *Gln1.2* genes expression among genotypes in root and shoot tissue

### Expression of *Gln2 gene*

OsGln2 expression was detected in both root and leaf tissues under all nitrogen treatments. In leaf tissues, expression was generally higher across the treatments than in root tissues (Fig. 3). Swarna showed high OsGln2 expression under ammonium sulfate and comparatively high expression across the evaluated treatments. In root tissues, the highest OsGln2 expression was observed under calcium nitrate in selected genotypes. Expression under the other treatments was moderate. Dagaddeshi showed relatively weak OsGln2 expression under some ammonium treatments (Fig 5).

**Fig 5.**
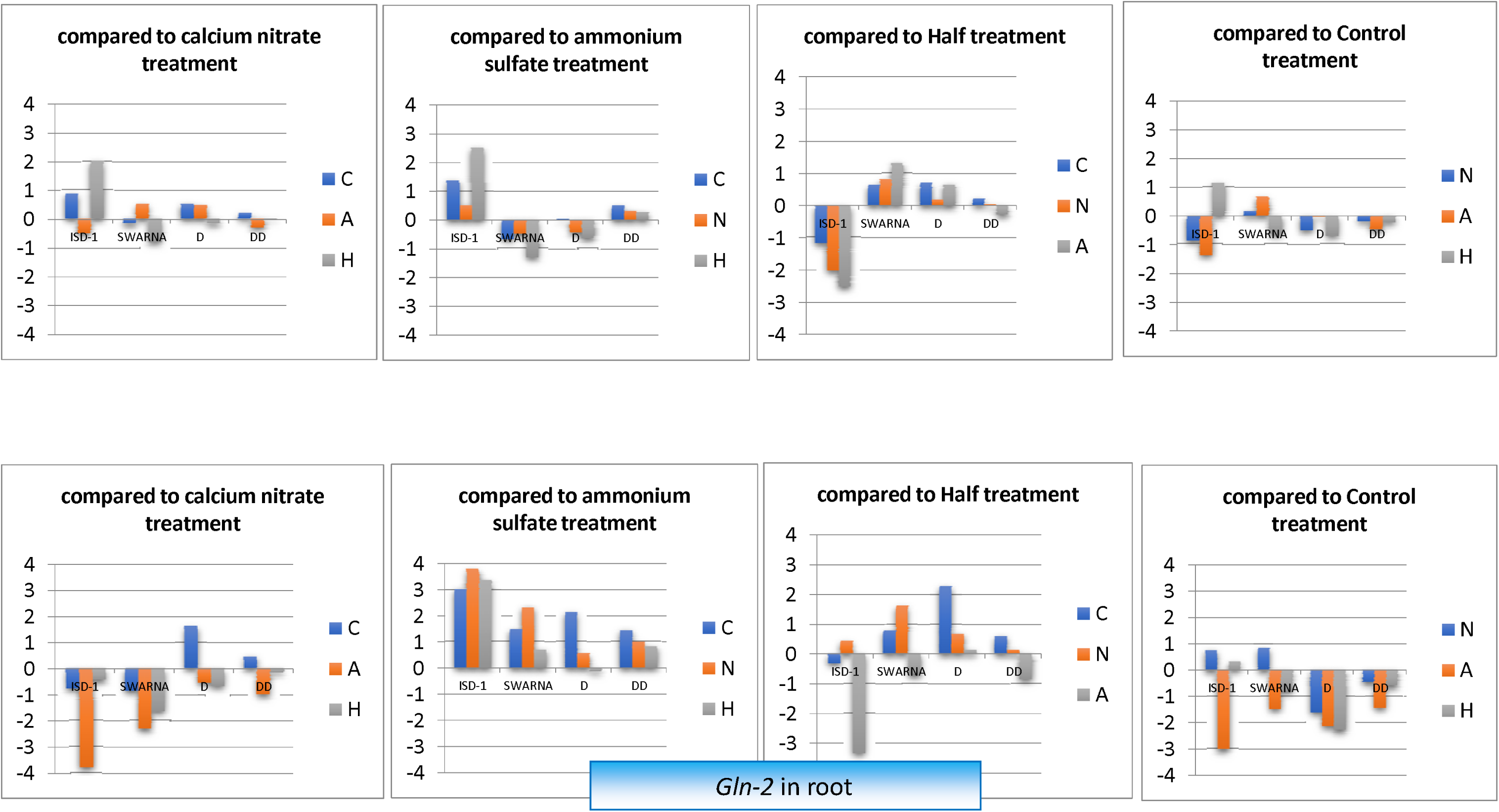
Comparison of treatments for *Gln2* genes expression among genotypes in root and shoot tissue

**Fig 6.**
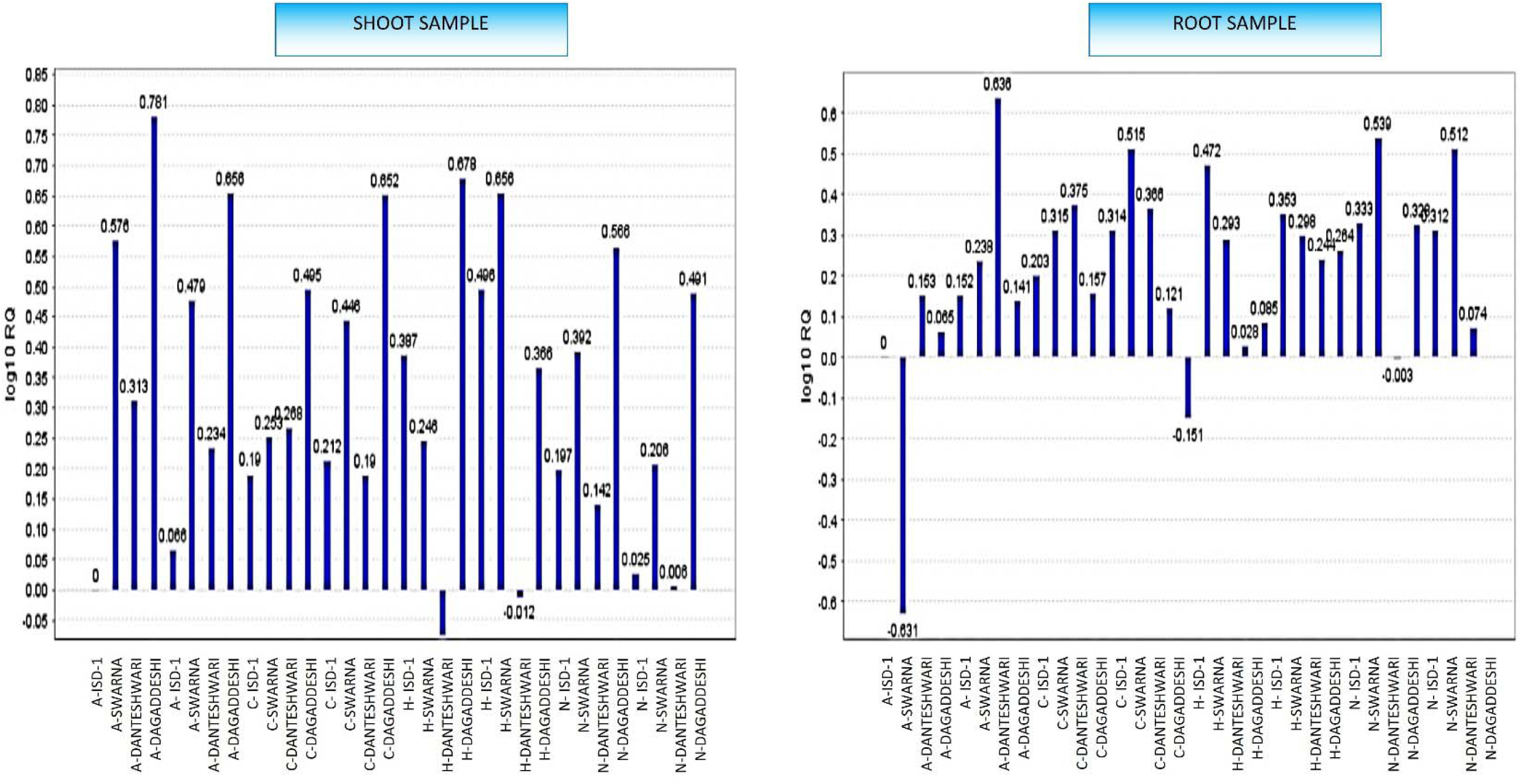
Expression profile of glutamate synthase gene in root and shoot under ammonium sulfate (A), AN [Ammonium Nitrate (C)], Half [half conc. of Ammonium Nitrate (H)] treatment and calcium nitrate (N) treatment

**Fig 7.**
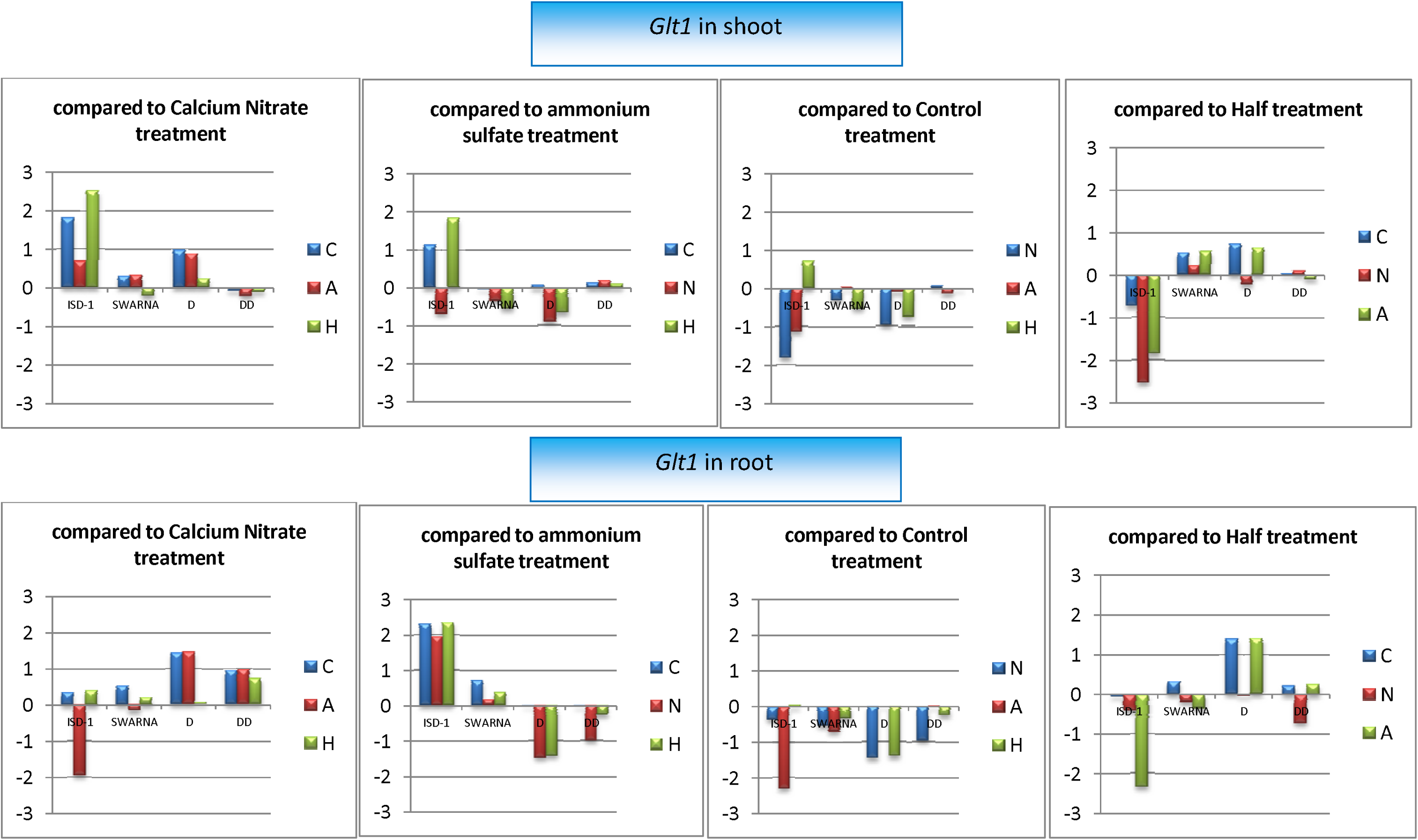
Comparison of treatments for *Glt1* genes expression among genotypes in root and shoot tissues

## Discussion

Rice production under direct-seeded and water-limited conditions requires genotypes that can maintain productivity while efficiently acquiring and assimilating nitrogen. In this study, substantial variation was observed among the 48 RILs for agronomic, physiological, biochemical, and molecular traits. The significant effects of genotype, nitrogen form, environment and their interactions demonstrate that nitrogen-use performance in rice is controlled by both genetic background and growing conditions. The results of genotype x nitrogen and genotype x environment interactions further indicate that rice genotypes do not respond uniformly to ammonium and nitrate, particularly under contrasting moisture regimes. Therefore, identifying nitrogen-efficient genotypes requires evaluation across both nutrient and water environments rather than under a single production condition.

Grain yield and biological yield were generally higher under ammonium nutrition than under nitrate nutrition, particularly in rainfed conditions. The response agrees with the physiological adaptation of rice to ammonium nutrition (Fan, Lu et al. 2025). Rice has traditionally evolved in flooded soils, where ammonium is commonly the predominant inorganic nitrogen form (Savant and De Datta 1982, Buresh, Ramesh Reddy et al. 2008). Ammonium can be assimilated directly through the GS/GOGAT pathway, whereas nitrate must first be transported into the plant and reduced to ammonium through the activities of nitrate reductase before it can be incorporated into amino acids. Consequently, ammonium assimilation requires fewer reduction steps and may impose a lower energetic cost on the plant than nitrate assimilation.

The superior performance of ammonium-treated plants under rainfed conditions is particularly important, as it contradicts the expectation that nitrate should be more readily available in aerobic, drained soils. Under aerobic conditions, ammonium may undergo nitrification to nitrate; however, the present results suggest that many genotypes in this population retained a greater capacity to absorb and utilize ammonium. This response may be related to the genetic background of the population and the relatively limited capacity of some genotypes to utilize nitrate efficiently. The result should therefore not be interpreted as evidence that ammonium is universally superior to nitrate in all rice production systems. Rather, it indicates that genotypic variation in nitrogen form preference can be exploited to develop varieties and fertilizer strategies for rainfed and direct-seeded rice.

The reduction in grain and biological yield under rainfed conditions confirms the strong influence of water availability on rice productivity. Water stress can restrict root growth, reduce nutrient mobility in the soil, impair nitrogen uptake, and limit photosynthesis and assimilate translocation to developing grains (Farooq, Wahid et al. 2009, Zhao, Wang et al. 2025). The significant reduction in yield under stress environments, therefore, reflects the combined effects of water limitation and reduced nitrogen acquisition. Nevertheless, several genotypes maintained relatively high yield across environments, indicating useful genetic variation for combined drought tolerance and nitrogen use. Ammonium nutrition also increased plant height, and, in several environments, early flowering. Increased tiller production is agronomically important because effective tillers contribute directly to panicle number and grain yield. The higher biological yield under ammonium treatment indicates that ammonium supported greater vegetative growth and biomass accumulation. However, greater biomass does not necessarily guarantee improved grain production; efficient partitioning of assimilates to the reproductive organs is also required. The variable harvest index observed among treatments indicates that nitrogen form influenced not only total biomass production but also the allocation of biomass between vegetative and reproductive structures. The relatively high harvest index under some control and nitrate treatments may reflect lower vegetative growth combined with a greater proportional allocation of available assimilates to grain, rather than superior overall productivity.

The strong positive correlation between biological and grain yield, especially under ammonium treatment, indicates that biomass production was an important contributor to yield in this population. This relationship was consistent under both irrigated and rainfed conditions, although it was weaker under rainfed nitrate conditions. The result suggests that ammonium-supported growth may provide a stronger base for grain formation, particularly when water availability is limited. Positive correlations between grain yield and flag leaf traits further emphasize the importance of maintaining an active photosynthetic canopy during the reproductive phase. Significant correlations involving flag leaf traits were treatment and environment-specific, suggesting that particular leaf traits may contribute to grain yield under certain nitrogen conditions.

The positive relationship between SPAD value and chlorophyll content confirms that SPAD measurements can provide a useful but indirect estimate of leaf nitrogen status in rice. SPAD readings were consistently higher under ammonium nutrition, whereas total chlorophyll showed a season and environment-dependent response. Since chlorophyll is directly associated with photosynthetic capacity, improved nitrogen status under ammonium nutrition may have contributed to greater biomass accumulation and grain yield. The relatively consistent relationship between SPAD and chlorophyll across environments also supports the use of SPAD readings for rapid field-level screening of nitrogen content. However, SPAD should be interpreted together with yield and NUE because a high chlorophyll concentration alone does not necessarily indicate efficient nitrogen conversion into grain.

The enzyme analysis data provides further support for the superior performance of ammonium nutrition. GS activity was highest under ammonium treatment, whereas GOGAT activity was highest under nitrate treatment. The enhanced GS activity under ammonium is physiologically plausible because ammonium is the direct substrate for GS (Liaw, Kuo et al. 1995). Higher GS activity would facilitate the rapid incorporation of ammonium into glutamine, thereby reducing the possibility of ammonium accumulation and toxicity while supplying nitrogen for amino acid, protein, chlorophyll and enzyme synthesis. In contrast, nitrate nutrition requires prior reduction to ammonium, which may explain the comparatively lower GS activity observed under nitrate treatment.

The higher GOGAT activity under nitrate nutrition suggests a different metabolic response. Following nitrate reduction, the resulting ammonium must be assimilated through the GS/GOGAT cycle. Increased GOGAT activity may therefore reflect a greater metabolic requirement for recycling glutamate and sustaining nitrogen assimilation when nitrate is supplied. The contrasting responses of GS and GOGAT indicate that these enzymes should not be interpreted independently. High GS activity may reflect efficient ammonium absorption, whereas high GOGAT activity under nitrate may indicate improved processing of nitrogen after nitrate reduction. The non-significant correlation between GOGAT activity and grain yield also suggests that high enzyme activity alone is not sufficient to predict agronomic performance.

In contrast, GS activity showed a positive but nonsignificant association with grain yield, and several high-yielding genotypes, including Swarna and IBD, expressed high GS activity under both ammonium and nitrate nutrition. This suggests a possible contribution of GS-mediated nitrogen assimilation to productivity; however, this relationship requires further validation. The strong GS activity observed in DxD 124-21 under ammonium and in 12.1(487)-237 under nitrate also demonstrates that different genotypes may possess distinct strategies to assimilate nitrogen. These lines could serve as useful donor materials or contrasting genotypes for breeding programs targeting nitrogen assimilation and adaptation to different forms of nitrogen. Nevertheless, GS activity should be considered a candidate biochemical indicator rather than a definitive marker until its relationship with NUE is validated across additional populations, environments, nitrogen rates and developmental stages.

The gene expression results partly supported the enzyme activity patterns, particularly for the stronger GS response under ammonium. However, OsGlt1 expression varied among genotypes, tissues and treatments and did not show a uniform pattern. Higher expression of Gln1.2 under ammonium sulfate and ammonium nitrate treatments indicates that this gene responds strongly to the availability of ammonium. The weaker expression under nitrate nutrition is consistent with previous reports showing that ammonium induces Gln1.2 expression more strongly than nitrate (Ishiyama, Inoue et al. 2004, Kojima, Minagawa et al. 2023). Higher Gln1.2 expression in high-performing genotypes such as Swarna and Danteshwari may indicate a greater capacity for ammonium assimilation and nitrogen remobilization. The expression of Gln2 in shoot tissues was relatively higher in selected genotypes under ammonium treatment and root tissues under nitrate treatment, suggesting tissue-specific regulation of nitrogen assimilation. Greater root expression under nitrate may reflect increased demand for nitrate uptake and assimilation, whereas greater shoot expression under ammonium may reflect active assimilation of ammonium in photosynthetic tissues.

The coordinated expression of Gln2 and Glt1 in Swarna supports the functional association between GS and GOGAT in the nitrogen assimilation pathway. However, the molecular data should be interpreted cautiously because gene expression was evaluated in only four genotypes under hydroponic conditions at the tillering stage under hydroponics. Expression patterns in young plants may not fully represent gene regulation during tillering, flowering, grain filling or drought stress in the field. In addition, increased transcript abundance does not always result in proportional increases in enzyme activity or nitrogen accumulation. Further studies should therefore combine gene expression with enzyme activity, tissue nitrogen concentration, nitrogen uptake and remobilization and grain nitrogen content.

Overall, ammonium produced higher mean biological yield, SPAD values, GS activity and grain yield under different experimental conditions, particularly in rainfed. However, chlorophyll responses and individual genotype responses varied with season, environment and form of nitrogen supplied. The study also identified genotypes with contrasting biochemical and molecular responses to different forms of nitrogen, including high GS activity in DxD124-21, 12.2(487), Swarna and IBD under selected treatments. These genotypes may provide useful material for further studies of nitrogen assimilation and acclimatization to water-limited environments. The higher GS activity and stronger expression of selected GS-related genes in some genotypes suggest a possible contribution of GS-mediated nitrogen assimilation. However, the nonsignificant GS-yield correlations indicate that this relationship requires further validation. However, significant genotype x nitrogen x environment interactions show that nitrogen efficiency is not a single, universally expressed trait. Breeding and nutrient management programs should therefore combine field performance with biochemical and molecular screening. Genotypes with strong GS activity under ammonium, enhanced GOGAT activity under nitrate, and stable yield under water limitation may provide valuable genetic resources for developing rice cultivars suited to direct-seeded and rainfed production systems.

## Conclusion

The findings indicate that the preference of forms of nitrogen in rice is not a fixed trait but depends on genotype and water regime. Ammonium nutrition provided a clear advantage for mean grain yield under rainfed conditions, but several genotypes performed better with nitrate, particularly under irrigated conditions. Therefore, fertilizer recommendations should consider both genotype and soil moisture conditions rather than assuming that one nitrogen form is universally superior. The contrasting GS and GOGAT responses further indicate that rice genotypes may use different biochemical strategies to assimilate ammonium and nitrate. Although selected genotypes showed promising enzyme activity and grain yield, these traits should be validated in additional environments and populations. The identified contrasting genotypes can serve as useful material for future breeding and nitrogen management studies targeting direct-seeded and water-limited rice production.

## Statement of Declaration

### Data availability

The data associated with this work are included in this manuscript

### Declaration of generative AI use

The authors report that generative AI was not used in their research or preparation of this manuscript.

### Author Contributions

Helan Baby Thomas carried out experiments, analyzed and compiled the data and wrote the manuscript; Toshy Agarwal helped with the biochemical and molecular work; Satish Verulkar is the group leader who conceptualized the idea and revised the manuscript.

